# The duration of mitosis and daughter cell size are modulated by nutrients in budding yeast

**DOI:** 10.1101/080648

**Authors:** Ricardo M. Leitao, Douglas R. Kellogg

## Abstract

The size of nearly all cells is modulated by nutrients. Thus, cells growing in poor nutrients can be nearly half the size of cells in rich nutrients. In budding yeast, cell size is thought to be controlled almost entirely by a mechanism that delays cell cycle entry until sufficient growth has occurred in G1 phase. Here, we show that most growth of a new daughter cell occurs in mitosis. When the rate of growth is slowed by poor nutrients, the duration of mitosis is increased, which suggests that cells compensate for slow growth in mitosis by increasing the duration of growth. The amount of growth required to complete mitosis is reduced in poor nutrients, leading to a large reduction in cell size. Together, these observations suggest that mechanisms that control the extent of growth in mitosis play a major role in cell size control in budding yeast.

## Introduction

Cell growth during the cell cycle must be precisely controlled to ensure that cell division yields two viable cells of a defined size. This is achieved by cell size checkpoints, which delay key cell cycle transitions until an appropriate amount of growth has occurred. The mechanisms by which cell size checkpoints measure growth and trigger cell cycle transitions are poorly understood.

An interesting feature of cell size checkpoints is that they can be modulated by nutrients. Thus, in many kinds of cells the amount of growth required to proceed through the cell cycle is reduced in poor nutrients, which can lead to a nearly 2-fold reduction in size (Young and Fantes, 1987; Johnston et al., 1977). Nutrient modulation of cell size is likely an adaptive response that allows cells to maximize the number of cell divisions that can occur when nutrients are limiting. Nutrient modulation of cell size is of interest because it likely works by modulating the threshold amount of growth required for cell cycle progression. Thus, discovering mechanisms of nutrient modulation of cell size should lead to broadly relevant insight into how cell size is controlled.

Cell size checkpoints are best understood in yeast, where two checkpoints have been defined. One operates at cell cycle entry in G1 phase, while the other operates at mitotic entry (Nurse, 1975; Johnston et al., 1977). The G1 phase checkpoint delays transcription of G1 cyclins, which is thought to be the critical event that marks commitment to enter the cell cycle (Cross, 1988; Nash et al., 1988). The mitotic entry checkpoint delays mitosis via the Wee1 kinase, which phosphorylates and inhibits mitotic Cdk1 (Nurse, 1975; Gould and Nurse, 1989).

In budding yeast, several lines of evidence suggest that cell size control occurs almost entirely at the G1 checkpoint. Budding yeast cell division is asymmetric, yielding a large mother cell and a small daughter cell. The small daughter cell spends more time undergoing growth in G1 prior to cell cycle entry (Johnston et al., 1977). This observation led to the initial idea of a G1 size checkpoint that blocks cell cycle entry until sufficient growth has occurred. The checkpoint is thought to control G1 cyclin transcription because loss of *CLN3,* the key early G1 cyclin that drives cell cycle entry, causes a delay in G1 phase (Cross, 1990). Cell growth continues during the delay, leading to abnormally large cells (Cross, 1988). Conversely, overexpression of *CLN3* causes cell cycle entry at a reduced cell size (Cross, 1988; Nash et al., 1988). In contrast, loss of the Wee1 kinase, a key component of the mitotic checkpoint, causes only mild cell size defects in budding yeast (Jorgensen, 2002; Harvey et al., 2005; Harvey and Kellogg, 2003). Together, these observations suggest that cell size control occurs primarily during G1.

Although significant cell size control occurs in G1 phase, there is evidence that important size control occurs at other phases in the cell cycle in budding yeast. For example, cells lacking all known regulators of the G1 cell size checkpoint show robust nutrient modulation of cell size (Jorgensen et al., 2004). This could be explained by the existence of additional G1 cell size control mechanisms that have yet to be discovered, but it could also suggest that normal nutrient modulation of cell size requires checkpoints that work outside of G1 phase. More evidence comes from the observation that daughter cells complete mitosis at a significantly smaller size in poor nutrients than in rich nutrients (Johnston et al., 1977). This suggests the existence of a checkpoint that operates after G1, during bud growth, to control the size at which daughter cells are born. This possibility has not received significant attention because early work suggested that the duration of daughter bud growth is invariant and independent of nutrients (Hartwell and Unger, 1977). As a result, it has been thought that birth of small daughter cells in poor nutrients is a simple consequence of their reduced growth rate, rather than active size control. However, this has not been tested by directly measuring the duration of daughter cell growth in rich and poor nutrients, so it remains possible that checkpoints actively modulate the extent of daughter cell growth to control cell size at completion of mitosis.

Further evidence for size control outside of G1 phase has come from analysis of nutrient modulation of cell size. Protein phosphatase 2A associated with the Rts1 regulatory subunit (PP2A^Rts1^) is required for nutrient modulation of cell size (Artiles et al., 2009). Proteome-wide analysis of proteins regulated by PP2A^Rts1^ revealed that it controls critical components of both the G1 phase and mitotic entry cell size checkpoints, as well as several key regulators of mitotic progression (Zapata et al., 2014). The fact that PP2A^Rts1^ is required for nutrient modulation of cell size, while regulators of the G1 checkpoint are not, could be explained by a model in which PP2A^Rts1^ controls mitotic cell size checkpoint mechanisms that play an important role in nutrient modulation of cell size.

Here, we set out to determine whether nutrient modulation of cell size occurs solely at the G1 checkpoint, or whether it also occurs at other times during the cell cycle. To do this, we investigated how nutrients affect cell growth, cell size and cell cycle progression throughout the cell cycle.

## Results and Discussion

### The duration of mitosis is modulated by nutrients

Previous work suggested that cell cycle events that occur after bud emergence have a constant duration that is independent of the growth rate set by nutrients (Hartwell and Unger, 1977). However, the limited tools available at the time meant that the duration of cell cycle events had to be inferred from indirect measurements. To more directly address this question, we first grew cells in a rich carbon source (2% dextrose) or a poor carbon source (2% glycerol + 2% ethanol) and determined the duration of mitosis by assaying levels of the mitotic cyclin Clb2 in synchronized cells. Clb2 persisted for a longer interval in cells growing in poor nutrients, which suggested that the duration of mitosis is increased (**Figure 1A**).

**Figure 1:**
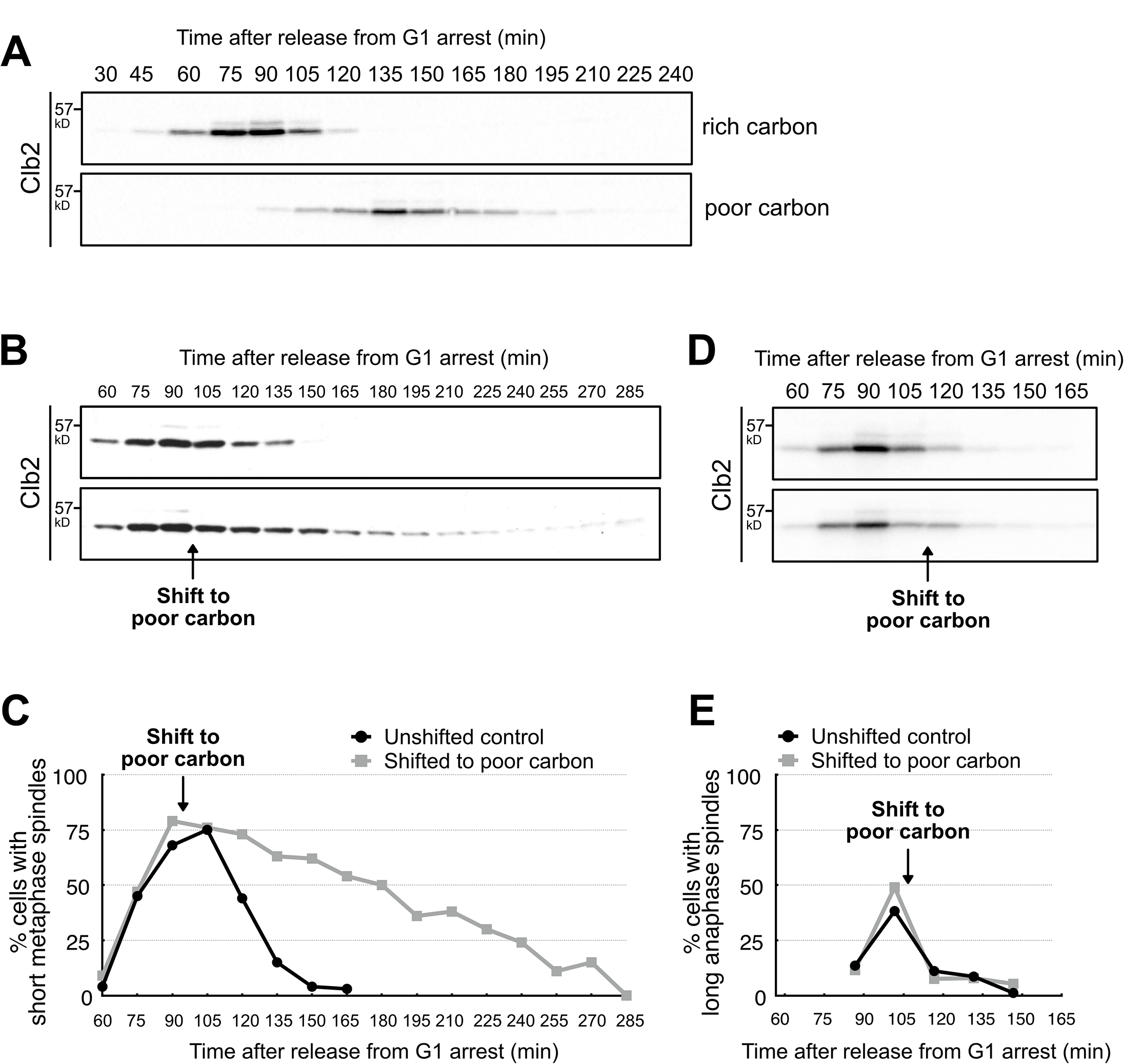
The duration of mitosis is modulated by nutrients. **(A)** Wild type cells growing in YPD (rich carbon) or YPG/E (poor carbon) were arrested in G1 phase by addition of mating pheromone. The cells were released from the arrest and levels of the mitotic cyclin Clb2 were assayed by western blot. **(B,C)** Cells growing in YPD were released from a G1 arrest. At 90 minutes, the culture was split and one half was washed into YPD and the other half was washed into YPG/E. Levels of the mitotic cyclin Clb2 were assayed by western blot **(B)** and cells with short metaphase spindles were assayed by immunofluorescence **(C)**. **(D,E)** Cells growing in YPD were released from a G1 arrest. At 105 minutes, the culture was split and one half was washed into YPD the other half was washed into YPG/E. Levels of the mitotic cyclin Clb2 were assayed by western blot **(D)** and long anaphase spindles were assayed by immunofluorescence **(E)**. Numbers on the left side of western blots indicate molecular weight in kilodaltons.

We next asked whether cells already in mitosis were sensitive to a shift from rich to poor carbon. Cells growing in rich carbon were synchronized and shifted to poor carbon when Clb2 reached peak levels and most cells had short mitotic spindles, indicating that they were in metaphase. A shift to poor carbon at this point in mitosis caused a prolonged metaphase delay, as well as delayed destruction of Clb2 (**Figures 1B,C**). In contrast, if cells were switched to poor carbon slightly later in mitosis, when cells were in anaphase, there was no delay in destruction of Clb2 or in completion of anaphase (**Figures 1D,E**). The insensitivity of anaphase cells to carbon source suggests that the metaphase delay is not due simply to a starvation response, which would likely affect both metaphase and anaphase.

To further investigate the effects of nutrients, we used fluorescence and bright field microscopy to simultaneously monitor daughter bud growth and mitotic events in living cells. Bud growth was monitored by plotting daughter bud volume as a function of time. To monitor key events of mitosis, mitotic spindle poles were marked with Spc42-GFP and the distance between poles was plotted as a function of time. Initiation of metaphase corresponds to the initial separation of spindle poles, while the duration of metaphase corresponds to the interval when spindle poles remain separated by 1-2 microns within the mother cell (Winey and O’Toole, 2001; Lianga et al., 2013). Initiation of anaphase is detected when spindle poles begin to move further apart and one pole migrates into the daughter cell. We defined the duration of anaphase as the interval between anaphase initiation and the time when the spindle poles reached their maximum distance apart. GFP-tagged myosin was used to detect completion of cytokinesis, which is marked by disappearance of the myosin ring (Lippincott and Li, 1998). In addition to the stages of mitosis, we defined an S/G2 interval as the time from bud emergence to spindle pole separation, and G1 phase as the time from when the daughter cell completes cytokinesis to the time that it initiates formation of a new daughter bud.

Bud growth and size were analyzed through a complete bud growth cycle, from the time the bud emerged from the mother cell to the time that it initiated formation of a new bud at the end of the next G1 phase. We utilized cells synchronized in G1 phase by mating pheromone arrest and release, as well as asynchronous cells. Cells synchronized by mating pheromone undergo growth during the arrest, and therefore initiate bud growth at a larger mother cell size than asynchronous cells (**Figure S1**). For both synchronous and asynchronous cells, there was not a statistically significant difference between average mother cell size in rich and poor carbon. This indicates that any differences in daughter cell growth or cell cycle timing between the two conditions are unlikely to be due to differences in mother cell size.

Representative data for synchronous cells growing in rich and poor carbon are shown in **Figures 2A** **and** **2B**, respectively. Examples of cell images taken during the course of bud growth are shown in **Figure 2C**. At least 25 cells were analyzed under each condition (See **Figures S2A,B for individual growth curves**).

**Figure 2:**
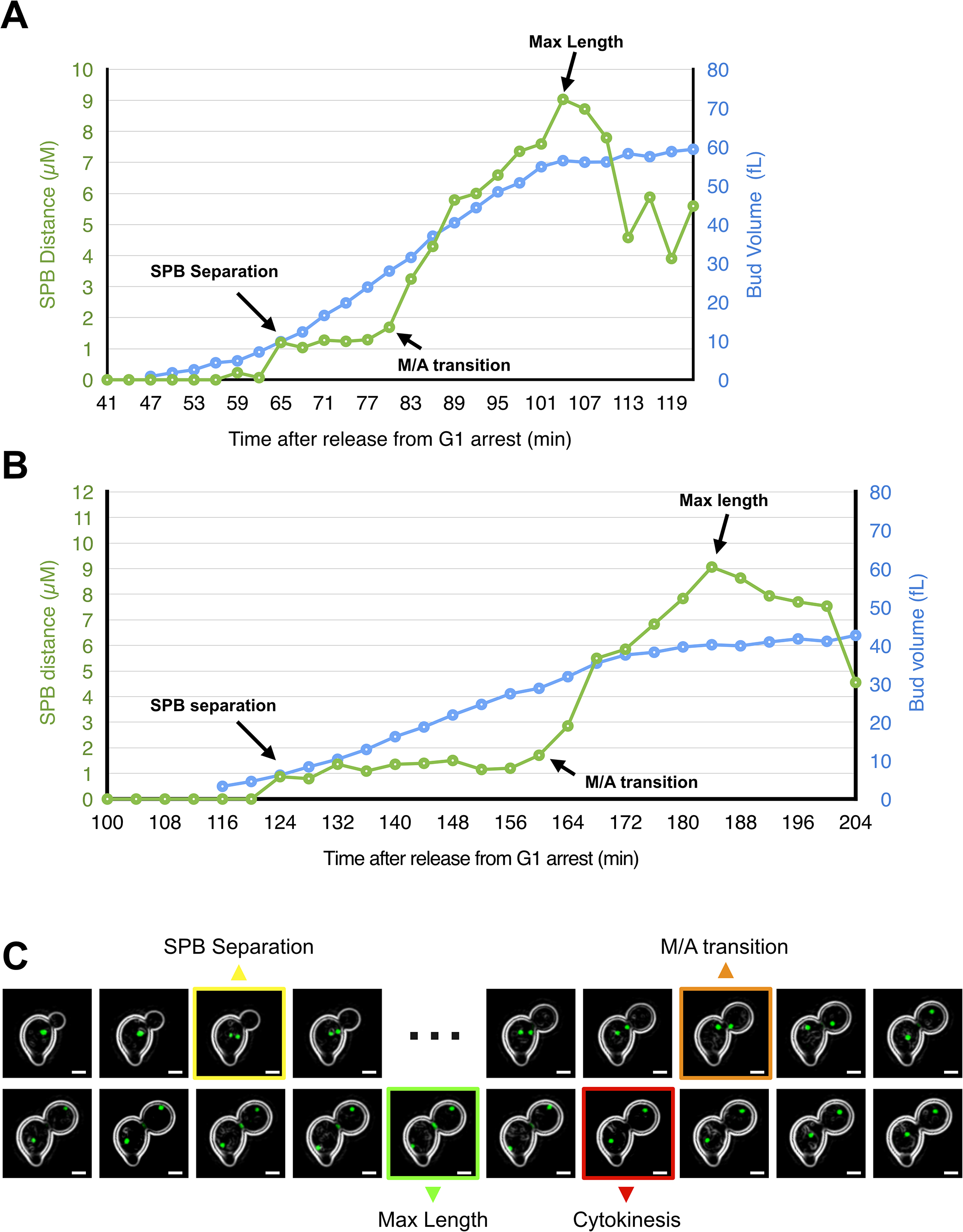
Simultaneous imaging of bud growth and mitotic spindle dynamics. **(A,B)** Representative growth curves for cells growing in rich carbon **(A)** or poor carbon **(B)**. The volume of the daughter bud is plotted in blue and the distance between spindle poles in green. **(C)** Contrast-enhanced images of a representative growing bud with GFP-tagged spindle poles (Spc42-GFP) and myosin ring (Myo1-GFP). Key transitions are highlighted. Images taken in metaphase that were omitted so that all key transition points could be shown are indicated with ▪▪▪. Scale bar indicates 2 uM.

The durations of all cell cycle stages comprising a complete bud growth cycle in rich and poor carbon are shown in **Figure 3A** for synchronous cells, and in **Figure 3B** for asynchronous cells (see **Figures S3A for scatter plots and p-values**). To focus on the effects of carbon source on mitosis, we also plotted separately the average durations of metaphase and anaphase for cells growing in rich or poor carbon (**Figures 3C,D**).

**Figure 3:**
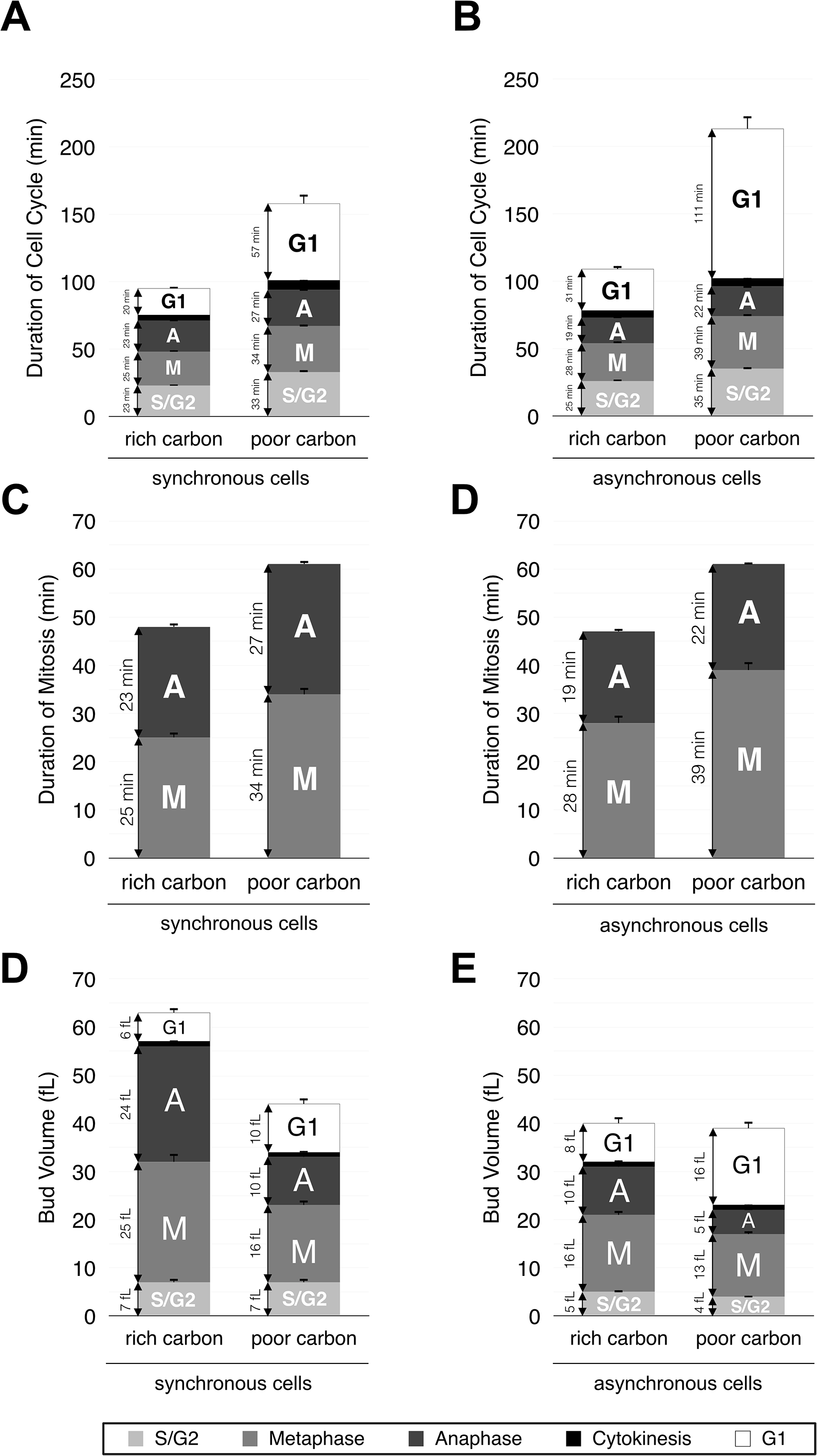
The duration of mitosis and cell size at completion of mitosis are modulated by nutrients. **(A,B)** Plots showing the average durations of all cell cycle stages for synchronous cells **(A)** or asynchronous cells **(B)** growing in rich or poor carbon. **(C,D)** Plots showing the average durations of metaphase and anaphase for synchronous cells **(A)** or asynchronous cells **(B)** growing in rich or poor carbon. **(E,F)** Plots showing the average growth in volume during all phases of the cell cycle for synchronous cells **(E)** or asynchronous cells **(F)** growing in rich or poor carbon. Error bars represent standard error of the mean.

The durations of both metaphase and anaphase were significantly increased in poor carbon. The overall average duration of mitosis was similar in the synchronous and asynchronous cells, although the fraction of mitosis spent in metaphase was slightly increased in asynchronous cells. The fraction of the growth cycle spent in G1 phase increased in poor carbon, as previously reported (Hartwell and Unger, 1977). The increase in G1 phase was greatest in asynchronous cells.

### Daughter cell size is modulated by nutrients

We next plotted bud size at completion of each cell cycle stage (**Figures 3E,F**; **see Figures S3B for scatter plots and p values**). Cells in poor carbon completed each stage of mitosis at a significantly smaller size. The data also show that in both rich and poor carbon, more growth in volume occurs during mitosis than in G1 phase. The increased size of daughter cells in synchronous cells relative to asynchronous cells is consistent with previous reports that mother cell size influences daughter cell size (Johnston et al., 1977; Schmoller et al., 2015).

The data are inconsistent with a model in which cells complete mitosis at a reduced size in poor nutrients simply because their growth rate is reduced, while the duration of mitotic events is unchanged. Rather, the duration of mitosis and cell size at completion of mitosis are modulated in response to changes in carbon source that cause large changes in growth rate. The data therefore suggest the existence of a nutrient-modulated mechanism that measures growth during mitosis and delays completion of mitosis until sufficient growth has occurred. This would explain why cells shifted from rich to poor carbon during metaphase undergo a prolonged mitotic delay, whereas cells shifted during anaphase do not (**Figure 1B-E**). If the delay were a consequence of a reduction in ATP or other metabolites needed for mitotic spindle events, one would expect to see delays in both metaphase and anaphase. Rather, we suggest that a shift to poor carbon during metaphase causes a delay because the daughter bud has not yet undergone sufficient growth, whereas a shift in anaphase does not cause a delay because buds have already reached the threshold amount of growth needed to complete mitosis in poor carbon. Since a large fraction of total growth occurs in mitosis, it would make sense that mechanisms that control the extent of growth in mitosis play a significant role in cell size control. The existence of major cell size control mechanisms in mitosis would explain why cells lacking critical regulators of the G1 size checkpoint still show robust nutrient modulation of cell size (Jorgensen et al., 2004). Work in fission yeast has suggested that there are mitotic cell size control mechanisms that act independently of Cdk1 inhibitory phosphorylation, which could explain why loss of Cdk1 inhibitory phosphorylation in budding yeast causes only modest effects on cell size (Wood and Nurse, 2013).

An alternative model is that the duration of mitosis is controlled by a nutrient modulated timer. In rich media, the timer would be set for a short duration of growth, while in poor media it would be set for a longer interval. However, comparison of the data from synchronous and asynchronous cells would appear to rule out a nutrient modulated timer model. Synchronous cells spend a total of 51 minutes in metaphase and anaphase in poor carbon, whereas asynchronous cells spend 61 minutes, despite growing under identical nutrient conditions. The difference is most likely due to a slower growth rate in asynchronous cells (see below), which would result in the need for a longer interval of growth to reach a critical amount of growth required for completion of mitosis. A timer model is also not consistent with the large variance in mitotic duration observed between individual cells growing under identical conditions (**Figure S3A**).

In both synchronous and asynchronous cells, the average size of mother cells initiating bud emergence was only slightly smaller in poor carbon compared to rich carbon, and the difference was not statistically significant (**Figure S1**). In addition, asynchronous daughter cells completed G1 phase at nearly identical sizes in rich and poor carbon (**Figure 3F**). These observations agree well with a previous study that found that cells growing in poor carbon complete late G1 phase at about 90% of the size of cells in rich carbon (Talia and Cross, 2007). This may at first seem paradoxical because poor carbon reduces average size nearly two-fold. However, cells growing in poor carbon are born at a dramatically reduced size, grow at a slower rate, and spend much more time in G1 phase. As a result, they spend more time at small sizes compared to cells in rich carbon, which contributes to a smaller average size when population averages are measured with a Coulter counter.

### The rate of growth is modulated during the cell cycle

Previous studies suggested that growth rate changes during the cell cycle, but did not include analysis of growth during specific stages of mitosis in unperturbed single cells (Goranov et al., 2009; Ferrezuelo et al., 2012). To extend these studies, we calculated average growth rates at each stage of the cell cycle in rich and poor carbon (**Figures 4A,B** **for synchronous cells and** **Figures 4C,D** **for asynchronous cells**). When the bud first emerges, growth is relatively slow. Entry into mitosis initiates a fast growing phase that lasts nearly the entire length of mitosis. As cells complete anaphase the growth rate slows. A slow rate of growth persists during G1 phase. Poor carbon reduced the rate of growth in mitosis by half, but caused smaller reductions in growth rate during other stages of the growth cycle. The rate of bud growth was greater in synchronized cells, which is due most likely to increased mother cell size (Schmoller et al., 2015). Previous studies have shown that polar bud growth is driven by Cdk1 activity; however, the signals that control bud growth at other stages of the cell cycle are unknown (McCusker et al., 2007). Moreover, the mechanisms and function of growth rate modulation during the cell cycle are unknown.

**Figure 4:**
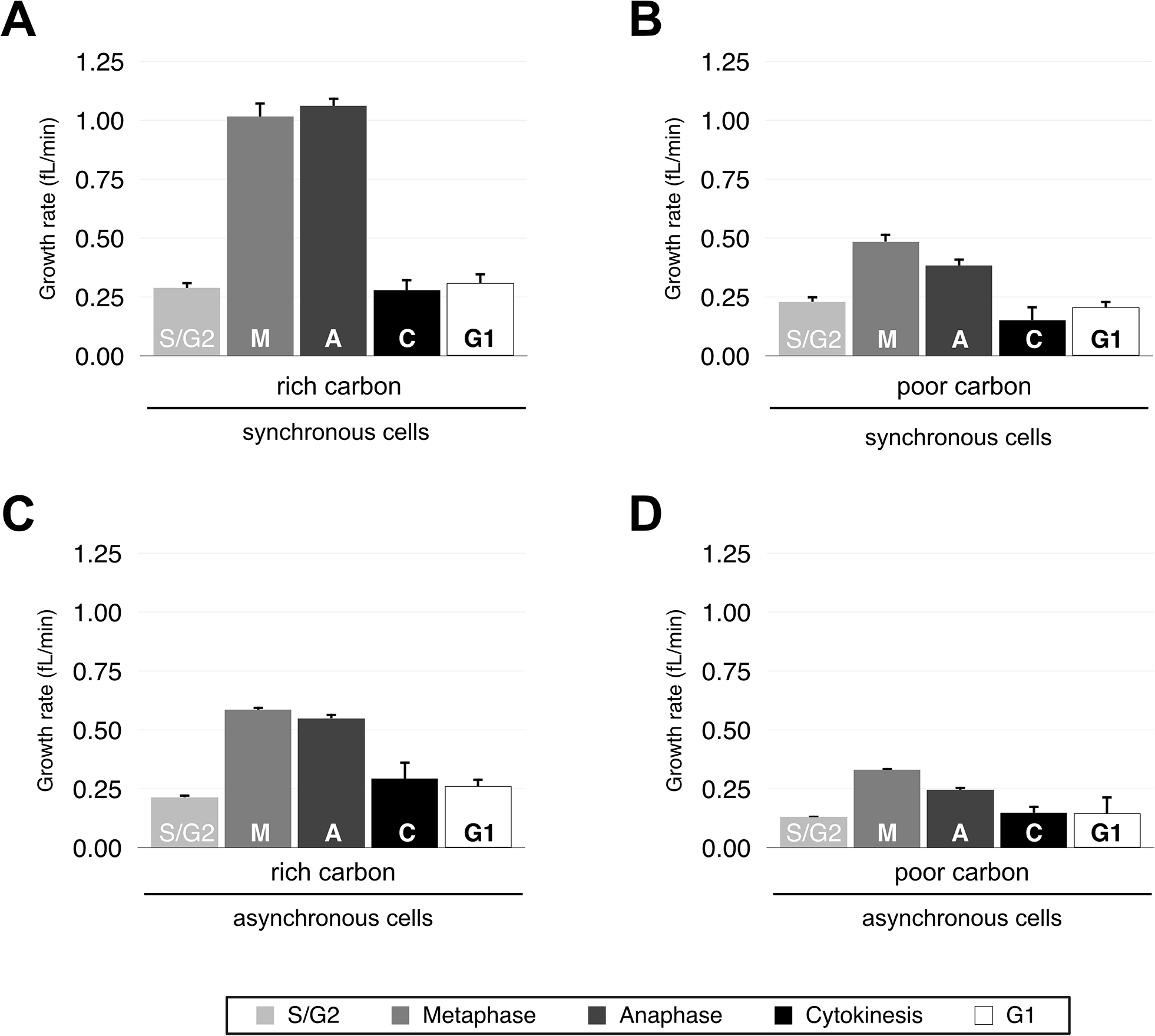
Cell growth rate is modulated during the cell cycle. The growth rate at each phase of the cell cycle was calculated as the average of individual cell growth rates. Growth rate was calculated by dividing the volume increase of the cell during a phase by the time the cell spent in that phase. **(A,B)** Data for synchronous cells growing in rich carbon **(A)** or poor carbon **(B)**. **(C,D)** Data for asynchronous cells growing in rich carbon **(C)** or poor carbon **(D)**. Error bars represent the standard error of the mean.

The discovery that most growth in volume occurs during a rapid growth phase in mitosis provides more evidence that cell size homeostasis requires tight control over the interval of mitotic growth. For example, a 20% change in the duration of growth in mitosis would have a large effect on cell size, whereas a 20% change in the duration of growth in G1 phase would have a smaller effect because the rate of growth in G1 phase is substantially slower.

### Effects of carbon source on daughter cell size are stronger than effects of mother cell size

Differences in cell growth and size between synchronous and asynchronous cells point to a strong influence of mother cell size on growth rate and daughter cell size. For example, synchronized cells initiate bud growth at a larger mother cell size compared to unsynchronized cells, and their daughter buds grow faster and complete mitosis at a larger size (**Figures S1, 3, 4**). Previous studies observed a similar correlation between mother cell size and daughter cell size (Johnston et al., 1977; Schmoller et al., 2015). These observations raised the possibility that the difference in daughter cell size at completion of mitosis in rich and poor carbon could be caused by differences in mother cell size due to nutrient modulation of cell size in G1 phase.

To further analyze the effects of mother cell size, we plotted the relationship between mother cell size and growth rate of the daughter bud in mitosis. Growth rate was positively correlated with mother cell size in both rich and poor carbon (**Figures 5A,B** **for synchronous and asynchronous cells, respectively**). Thus, daughters of large mothers grew faster than daughters of small mothers, consistent with the idea that mother cell size influences biosynthetic capacity. However, carbon source had a stronger influence on growth rate than mother cell size. This can be seen by the fact that mothers of similar size in rich and poor carbon had daughter buds that grew at different rates, which was true across the entire range of mother cell sizes.

**Figure 5:**
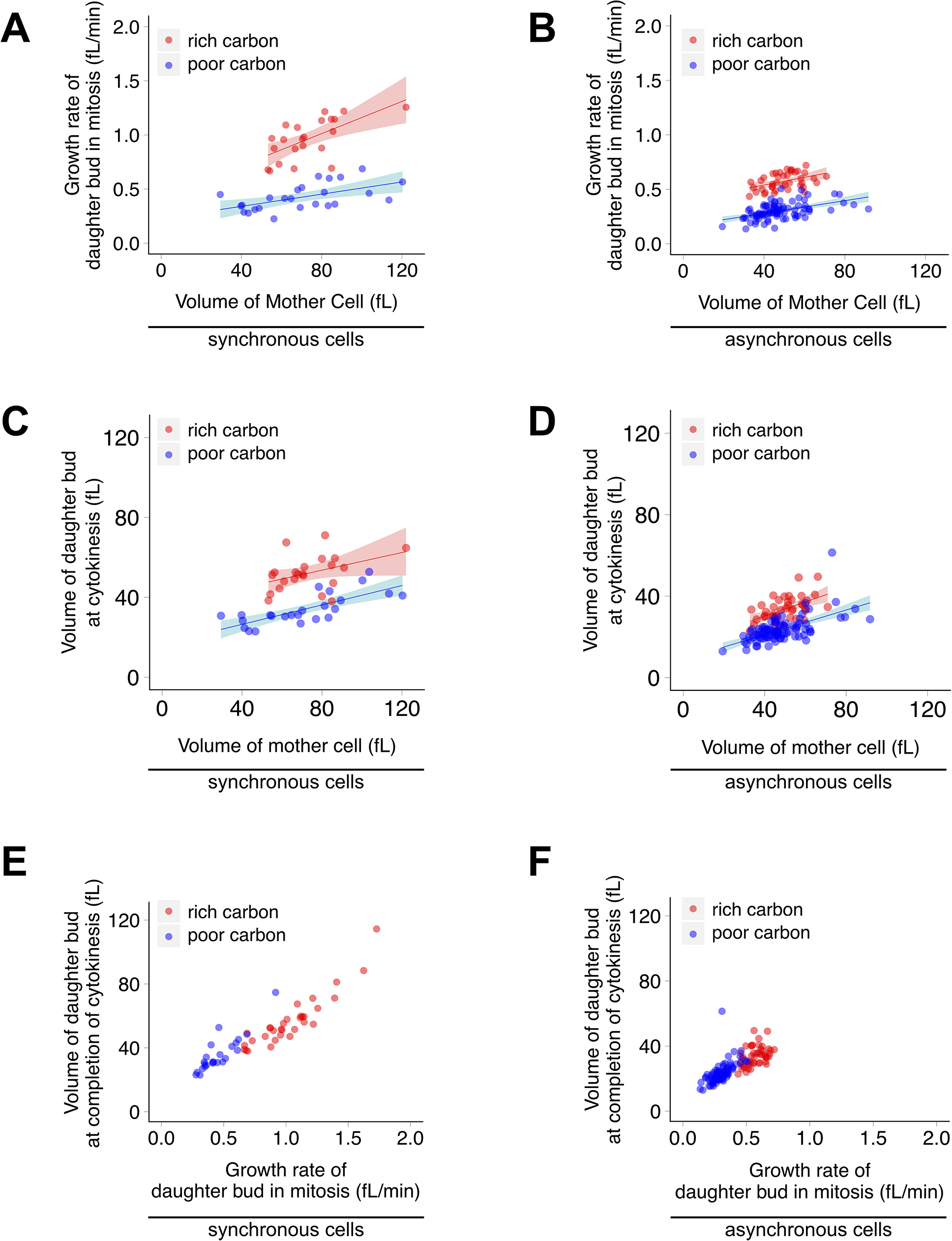
Cell size at completion of cytokinesis is proportional to the growth rate during mitosis. **(A,B)** The growth rate in mitosis of each daughter bud was plotted against the volume of its mother cell for synchronous cells **(A)** and asynchronous cells **(B)**. **(C,D)** The volume of each daughter bud at cytokinesis was plotted against the size of its mother for synchronous cells **(C)** and asynchronous cells **(D)**. **(E,F)** The volume of daughter cells at cytokinesis was plotted against their growth rate during mitosis for synchronous cells **(E)**, and asynchronous cells **(F)**. Red dots represent cells in rich carbon. Blue dots represent cells in poor carbon. Smooth lines are logistic regressions of the data. Shaded areas represent 95% confidence interval.

We also plotted daughter cell size at completion of cytokinesis versus mother cell size (**Figures 5C,D**). Daughter cell size was positively correlated with mother cell size in both conditions. However, the influence of carbon source was again much stronger. Mother cells growing in poor carbon that were the same size as mother cells in rich carbon consistently produced much smaller daughter cells. Together, these data indicate that effects of carbon source on daughter cell size can not be due solely to differences in mother cell size.

The effects of mother cell size could explain why synchronized cells in rich carbon completed late G1 phase at a larger size than their counterparts in poor carbon (**Figure 3E**). Note that synchronized cells in both rich and poor carbon appear to overshoot the size at which asynchronous cells complete late G1 phase, which could be due to increased mother cell size in the synchronized cells. The large mothers in synchronized cells in rich carbon drive a high rate of growth, which could lead to greater overshooting of the threshold amount of growth required for G1 progression. Asynchronous cells in both rich and poor carbon have smaller mother cells and are born at smaller sizes relative to synchronized cells. In this case, compensatory growth in G1 appears to become more important to bring the daughter cell up to a minimal threshold size before cell cycle entry.

### Cell size at completion of mitosis is correlated with growth rate during mitosis

Previous studies found that cell size at the end of G1 phase is correlated with growth rate during G1 phase (Ferrezuelo et al., 2012; Johnston et al., 1979). The correlation holds true when comparing cells growing in the same carbon source and when comparing cells growing in different carbon sources. To determine whether a similar relationship exists for growth during mitosis, we plotted daughter cell size at cytokinesis as a function of daughter bud growth rate during mitosis for cells growing in rich or poor carbon (**Figure 5E,F**). Daughter cell size was positively correlated with growth rate under both conditions. Thus, cell size at all key cell cycle transitions is correlated with growth rate.

Because cell size is proportional to growth rate, faster growing cells should always give rise to larger daughter cells. Moreover, since growth rate is proportional to size, larger mother cells should have a higher growth rate, leading to ever larger daughter cells. In this case, what limits cell size? The plot of daughter bud size at cytokinesis as a function of mother cell size revealed that daughter cell size indeed increases with mother cell size, but the ratio of mother size to daughter size is not constant across the range of mother cell sizes (**Figures 5C,D**). In other words, small mothers produce daughters of nearly equal size, while very large mothers produce daughters that are nearly half the size of the mother (Johnston et al., 1977). This relationship would correct large variations in mother cell size, which could be the result of growth during cell cycle delays induced by other checkpoints, such as the spindle checkpoint or DNA damage checkpoints.

The strong correlation between growth rate and cell size is difficult to reconcile with simple cell size checkpoint models in which a threshold volume must be reached to pass the checkpoint. If a specific volume must be reached to pass a checkpoint, the rate at which the cell reaches that volume should not influence the final volume at which the cell passes the checkpoint. One way to reconcile the idea of a set threshold volume with growth rate dependence would be to imagine that cell size checkpoint thresholds are noisy and imperfect.

In this view, faster growing cells will overshoot the threshold size more than slow growing cells, leading to increased size. However, this model would not explain nutrient modulation of cell size. Thus, another model could be that cells measure their growth rate and set cell size thresholds to match growth rate (Jorgensen et al., 2004). Alternatively, the same signals that set the growth rate could also set the cell size threshold. Both models would explain nutrient modulation of cell size, since nutrients modulate growth rate.

## Materials and Methods

### Yeast strains and media

The genotype of the strain used in this study is *SPC42-GFP::HIS3 MYO1-GFP::TRP1 leu2-3,112 ura3-1 can1-100 ade2-1 his3-11,15 trp1-1 GAL+, ssd1-d2* (W303 background). Genetic alterations were carried out using one-step PCR-based integration at the endogenous locus (Longtine et al., 1998) or by genetic crossing.

For cell cycle time courses cells were grown in YP media (1% yeast extract, 2% peptone, 8ml/L adenine) supplemented with 2% dextrose (YPD), or with 2% glycerol and 2% ethanol (YPG/E). For microscopy, cells were grown in complete synthetic media (CSM) supplemented with 2% dextrose (CSM-Dex) or 2% glycerol and 2% ethanol (CSM-G/E).

### Microscopy

Cells were grown overnight in CSM-DEX or CSM-G/E at room temperature with constant rotation to an optical density near 0.1 at λ600. 5 ml of culture were arrested with α factor at 0.5 μg/ml for 3-4 hours. Cells were released from the arrest by 3 consecutive washes with the same media and re-suspended in 500 μl of media. Approximately 200 μl of cell suspension were spotted on a concanavalin A-treated glass bottom dish with a 10 mm micro-well #1.5 cover glass. Cells were adhered for 5 min and unbound cells were washed away by repeated washing with 1 ml pre-warmed media. The dish was then flooded with 3 ml of media and placed on a temperature-controlled microscope stage set to 27°C (Pecon Tempcontrol 37-2 digital). The temperature of the media was monitored throughout the experiment using a MicroTemp TQ1 reader coupled to a Teflon insulated K-type thermocouple (Omega). The probe was placed in contact with the glass bottom near the contact area between the objective and the dish. Temperature was maintained at 27 ^+^/_-_ 1°C; imaging sessions where the temperature varied beyond this limit were rejected from final analysis. Brightfield and fluorescent images were acquired simultaneously using a Zeiss LSM 5 Pascal **Axiovert 200M inverted** microscope and a Plan-Apochromat 63x/1.4 oil objective. 488 nm light was obtained from an argon laser light source using a (488/543/633) primary dichroic beam splitter (HFT). The laser was set to 0.7% intensity. For green fluorescence images, light was collected through a long pass 505 emission filter using a 1 AU size pinhole. Brightfield images were collected using the transmitted light detector. Optical sections were taken for a total of 11 z-planes every 0.5 μm with frame averaging set to 2, to reduce noise. The total exposure was kept as low as possible to reduce photo-damage (1.60 μs dwell time per pixel, image dimension set to 512 x 512 px, and pixel size set to 0.14 x 0.14 μm). Images were acquired at 3 min intervals for rich carbon and 4 min intervals for poor carbon and recorded via the Zen 2000 interface.

### Image analysis

Image analysis was performed using ImageJ (Schneider et al., 2012; Linkert et al., 2010). The ImageJ plug-ins StackReg and MultiStackReg were used for post-acquisition image-stabilization (Thevenaz et al., 1998; Busse). Stabilized bright-field images were processed using the imageJ plug-in FindFocusedSlices and the volume of growing buds was determined using BudJ (Ferrezuelo et al., 2012; Tseng). Bud volumes were measured for buds whose focal plane was no more than 1.5 μm away (3 z-steps) from their mother’s focal plane. Sum projections of z-stacks were treated using a 2 px mean filter, and brightness/contrast levels were adjusted to reduce background noise. The treated pseudo-colored green fluorescent images were overlapped with the outlines of the imaged cells for reference (outlines were generated using the “find edges” command over a sum projection of all z-stacks of bright field images). Positions of spindle poles (SPBs) were determined using the crosshair tool (set to auto-measure and auto-next) and distance between the 2 SPBs was determined using the mathematical formula for the distance between 2 points. Disappearance of the Myo1 ring was determined empirically by observation of GFP signal at the bud neck.

### Statistical analysis

Data acquired from ImageJ was analyzed using Apple Numbers, R (The R Core Team, 2016), RStudio (RStudio Team, 2015), and the R package ggplot2 (Wickham, 2009). p-values were calculated using a Welch Two Sample t-test and a 95% confidence interval.

### Cell cycle time courses and western blotting

For western blot time courses, cells were grown overnight at room temperature in liquid YPD or YPG/E to an optical density of 0.5 at λ600. Because optical density is affected by cell size, we normalized cell numbers by counting cells with a Coulter counter (Beckman Coulter) when comparing cells grown in rich or poor carbon.

G1 synchronization was achieved by arresting cells with α factor at a concentration of 0.5 μg/ml until at least 90% of cells were unbudded. Cells were released from the arrest by washing 3× with fresh media. Time courses were performed at 25°C with constant agitation. To prevent cells from re-entering the following cell cycle α factor was added back after most cells had budded. For nutrient shift time courses, a single culture was arrested and split at the moment of release from the G1 arrest. At the time of the shift both cultures were washed 3× with room temperature YPD (control) or YPG/E (shifted cells) by centrifugation for 1 min at 3,000 × g (Eppendorf 5702). The volume of the culture was restored to its original volume prior to the washes and cultures were placed back at 25°C.

For western blots, 1.6 ml samples were collected at regular intervals, pelleted and flash frozen in presence of 200 μl of glass beads. Cell lysis and western blotting were carried out as previously described (Harvey et al., 2005).

Immunofluorescence analysis of mitotic spindles was carried out as previously described (Pringle et al., 1991). Multi-plane imaging of slides was performed using a Leica DM5500 B Widefield Microscope and a 63x/ 0.6-1.4 oil objective. Spindles were counted in ImageJ. Data was processed and plotted using Apple Numbers.

## Supplemental Material

Figure S1 shows distributions of mother cell volumes for cells growing in rich or poor carbon and for synchronous and asynchronous cultures. Black horizontal lines and adjacent numbers represent the average value for the data in each dot plot. The significance of the difference between two conditions is given as p-values above each plot. Figure S2 shows growth curve data for cells growing in rich or poor carbon. Black represents S/G2 phase. Blue represents metaphase. Pink represents anaphase. Yellow represents G1 phase. Curves where yellow is absent are from cells that could not be followed through a complete G1 phase or where the timing of the appearance of the daughter bud could not be determined with confidence. All curves have the same Y-axis scale (0-125 fL). The X-axis scale is variable between curves. Panel S2A shows growth curves for cells growing in rich carbon. Curves for all 32 measured cells are shown. Panel S2B shows growth curves for cells growing in poor carbon. Several curves were omitted because they yielded clear data for one stage of mitosis, but not for others, due to imaging limitations. Data from these curves were used if they yielded high confidence data for one of the mitotic stages. Figure S3 shows dot plot versions of the data used to generate Figure 3. Black horizontal lines and adjacent numbers represent the average value for the data in each dot plot. The significance of the difference between two conditions is given as p-values above each plot. Panel S3A shows dot plot versions of the data used to generate Figures 3A,B. Panel S3B shows dot plot versions of the data used to generate Figures 3C,D.

## Acknowledgments

We thank Ben Abrams, Facilities Manager for the UCSC Life Sciences Microscopy Center for support and mentoring with all microscopy related techniques, and the Aldea lab for sharing BudJ: an ImageJ plugin to analyze images of budding yeast cells (http://www.ibmb.csic.es/home/maldea). R.L. was supported by a fellowship from the “Fundação para a Ciência e a Tecnologia” (FCT), with funds from “Programa Operacional Potencial Humano/ Fundo Social Europeu” (POPH/FSE), under the fellowship SFRH/BD/75004/2010. This work was supported by National Institutes of Health Grant R01- GM053959.

The authors declare no competing interests.

## Author Contributions

Ricardo Leitao contributed conceptualization, investigation, data curation, formal analysis, methodology, software, visualization, writing, review, and editing. Douglas Kellogg contributed conceptualization, methodology, writing, review, editing, supervision, project administration and funding acquisition.

---

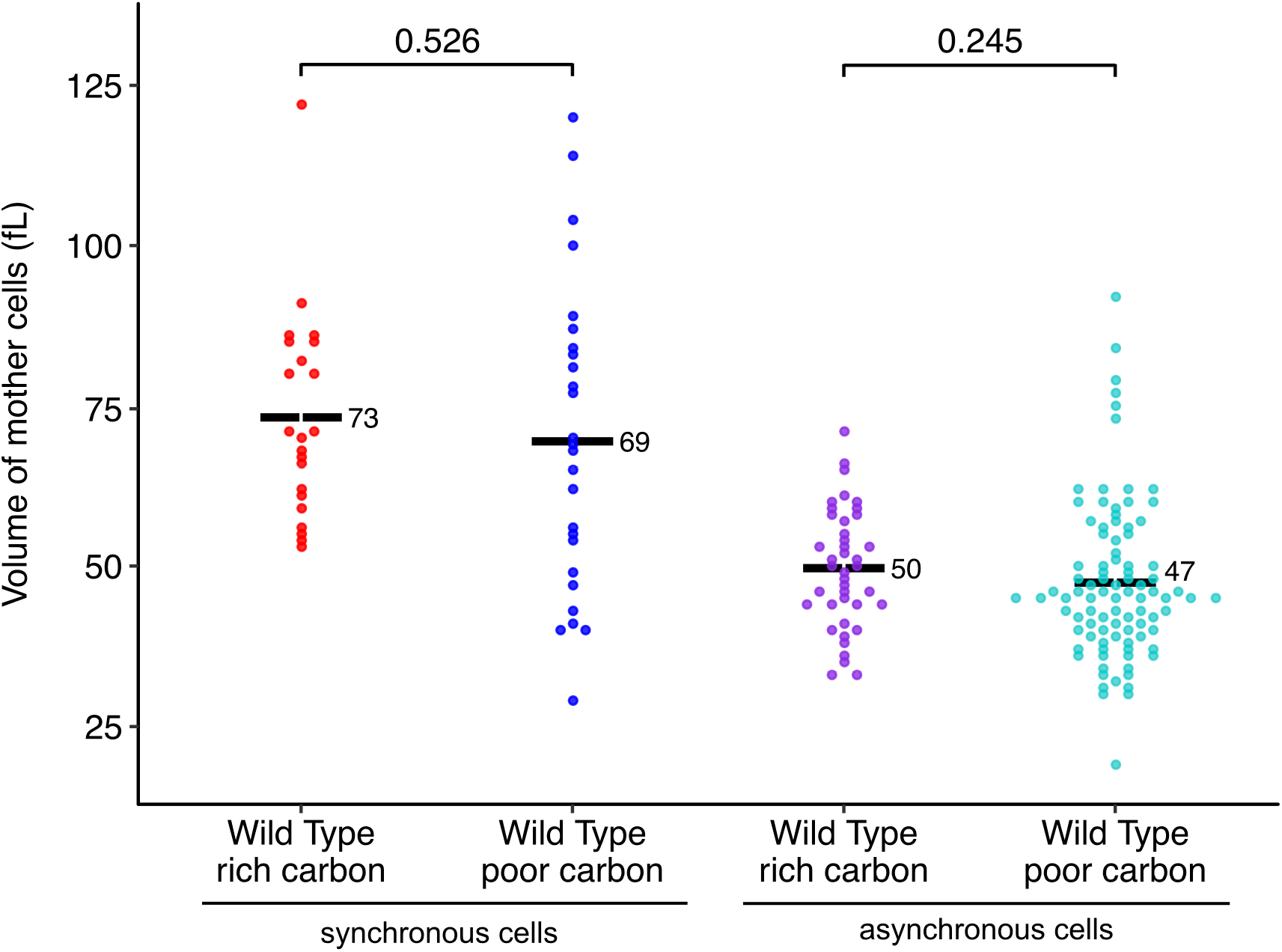
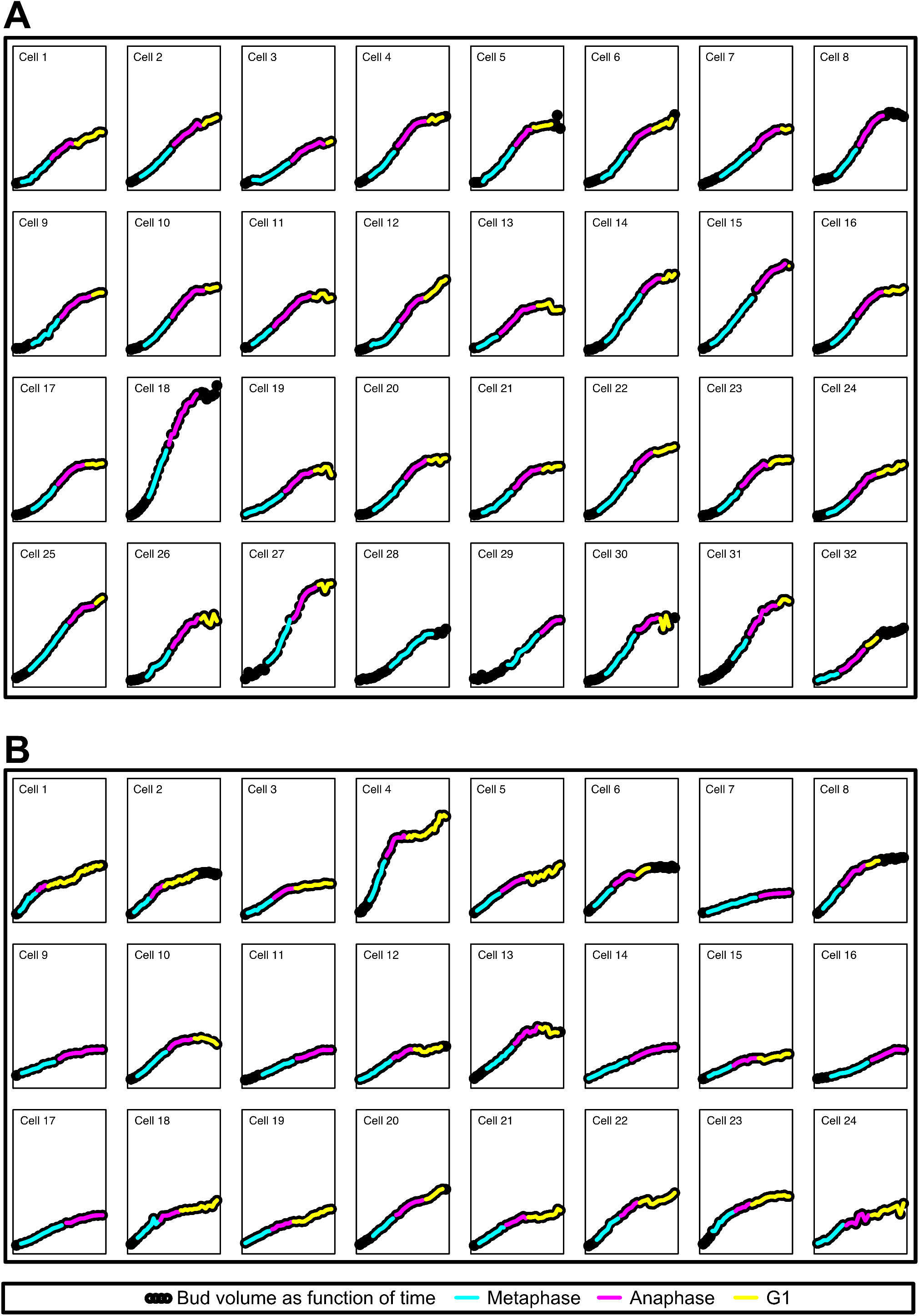
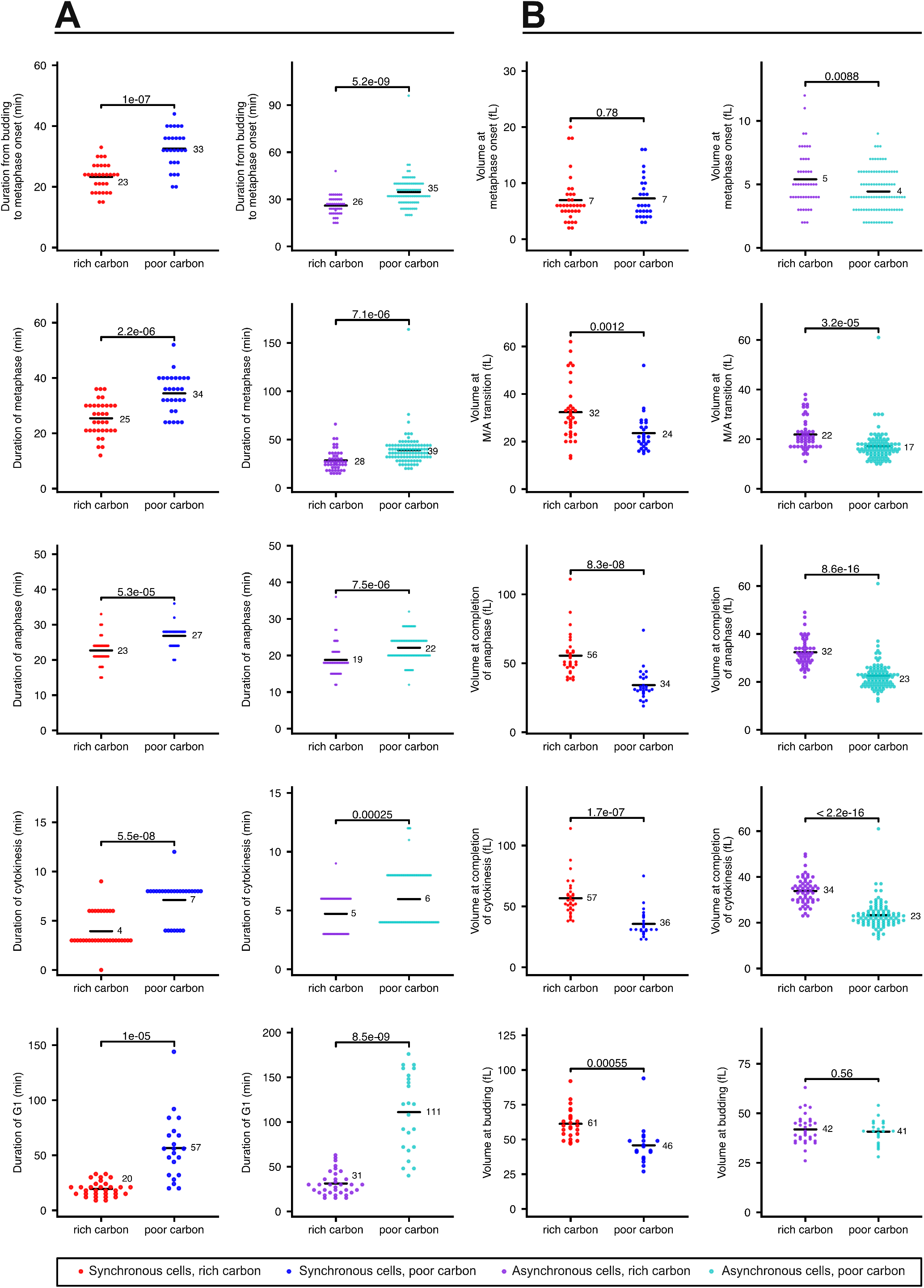

## References

Artiles, K., S.D. Anastasia, D. McCusker, and D.R. Kellogg. 2009. The Rts1 Regulatory Subunit of Protein Phosphatase 2A Is Required for Control of G1 Cyclin Transcription and Nutrient Modulation of Cell Size. PLoS Genet. 5:e1000727 EP –. doi:10.1371/journal.pgen.1000727.

Busse, B. (n.d.). MultiStackReg V1.45. http://bradbusse.net/downloads.html.

Cross, F.R. 1988. DAF1, a mutant gene affecting size control, pheromone arrest, and cell cycle kinetics of Saccharomyces cerevisiae. Mol. Cell. Biol. 8:4675–4684. doi:10.1128/MCB.8.11.4675.

Cross, F.R. 1990. Cell cycle arrest caused by CLN gene deficiency in Saccharomyces cerevisiae resembles START-I arrest and is independent of the mating-pheromone signalling pathway. Mol. Cell. Biol. 10:6482–6490. doi:10.1128/MCB.10.12.6482.

Ferrezuelo, F., N. Colomina, A. Palmisano, E. Garí, C. Gallego, A. Csikász-Nagy, and M. Aldea. 2012. The critical size is set at a single-cell level by growth rate to attain homeostasis and adaptation. Nat Commun. 3:1012–11. doi:10.1038/ncomms2015.

Goranov, A.I., M. Cook, M. Ricicova, G. Ben-Ari, C. Gonzalez, C. Hansen, M. Tyers, and A. Amon. 2009. The rate of cell growth is governed by cell cycle stage. Genes & Development. 23:1408–1422. doi:10.1101/gad.1777309.

Gould, K.L., and P. Nurse. 1989. Tyrosine phosphorylation of the fission yeast cdc2+ protein kinase regulates entry into mitosis. Nature. 342:39–45. doi:10.1038/342039a0.

Hartwell, L.H., and M.W. Unger. 1977. Unequal division in Saccharomyces cerevisiae and its implications for the control of cell division. The Journal of Cell Biology. 75:422–435. doi:10.1083/jcb.75.2.422.

Harvey, S.L., A. Charlet, W. Haas, S.P. Gygi, and D.R. Kellogg. 2005. Cdk1-dependent regulation of the mitotic inhibitor Wee1. Cell. 122:407–420. doi:10.1016/j.cell.2005.05.029.

Harvey, S.L., and D.R. Kellogg. 2003. Conservation of Mechanisms Controlling Entry into Mitosis: Budding Yeast Wee1 Delays Entry into Mitosis and Is Required for Cell Size Control. Current Biology. 13:264–275. doi:10.1016/S0960-9822(03)00049-6.

Johnston, G.C., C.W. Ehrhardt, A. Lorincz, and B.L.A. Carter. 1979. Regulation of cell size in the yeast Saccharomyces cerevisiae. J. Bacteriol. 137:1–5.

Johnston, G.C., J.R. Pringle, and L.H. Hartwell. 1977. Coordination of growth with cell division in the yeast Saccharomyces cerevisiae. Exp. Cell Res. 105:79–98. doi:doi: 10.1016/0014- 4827(77)90154-9.

Jorgensen, P. 2002. Systematic Identification of Pathways That Couple Cell Growth and Division in Yeast. Science. 297:395–400. doi:10.1126/science.1070850.

Jorgensen, P., I. Rupes, J.R. Sharom, L. Schneper, J.R. Broach, and M. Tyers. 2004. A dynamic transcriptional network communicates growth potential to ribosome synthesis and critical cell size. Genes & Development. 18:2491–2505. doi:10.1101/gad.1228804.

Lippincott, J., and R. Li. 1998. Sequential assembly of myosin II, an IQGAP-like protein, and filamentous actin to a ring structure involved in budding yeast cytokinesis. J. Cell Biol. 140:355–366. doi:10.1083/jcb.140.2.355.

Longtine, M.S., A. McKenzie, D.J. Demarini, N.G. Shah, A. Wach, A. Brachat, P. Philippsen, and J.R. Pringle. 1998. Additional modules for versatile and economical PCR-based gene deletion and modification in Saccharomyces cerevisiae. Yeast. 14:953–961. doi:10.1002/(SICI)1097-0061(199807)14:10<953::AID-YEA293>3.0.CO;2-U.

McCusker, D., C. Denison, S. Anderson, T.A. Egelhofer, J.R.I. Yates, S.P. Gygi, and D.R. Kellogg. 2007. Cdk1 coordinates cell-surface growth with the cell cycle. Nat Cell Biol. 9:506–U45. doi:10.1038/ncb1568.

Nash, R., G. Tokiwa, S. Anand, K. Erickson, and A.B. Futcher. 1988. The WHI1+ gene of Saccharomyces cerevisiae tethers cell division to cell size and is a cyclin homolog. EMBO J. 7:4335–4346.

Nurse, P. 1975. Genetic control of cell size at cell division in yeast. Nature. 256:547–551. doi: 10.1038/256547a0.

Pringle, J.R., A.E.M. Adams, D.G. Drubin, and B.K. Haarer. 1991. Immunofluorescence methods for yeast. In Methods in Enzymology. Academic Press. 565–602.

RStudio Team. 2015. RStudio: Integrated Development Environment for R. http://www.rstudio.com.

Schmoller, K.M., J.J. Turner, M. Kõivomägi, and J.M. Skotheim. 2015. Dilution of the cell cycle inhibitor Whi5 controls budding-yeast cell size. Nature. 526:268–272. doi: 10.1038/nature14908.

The R Core Team. 2016. R: A language and environment for statistical computing.

Thevenaz, P., U.E. Ruttimann, and M. Unser. 1998. A pyramid approach to subpixel registration based on intensity. IEEE Transactions on Image Processing. 7:27–41. doi: 10.1109/83.650848.

Tseng, Q. Find Focused Slices. https://sites.google.com/site/qingzongtseng/find-focus.

Wickham, H. 2009. ggplot2: Elegant Graphics for Data Analysis. Springer-Verlag, New York.

Winey, M., and E. O’Toole. 2001. The spindle cycle in budding yeast. Nat Cell Biol. 3:E23–E27. doi: 10.1038/35050663.

Wood, E., and P. Nurse. 2013. Pom1 and cell size homeostasis in fission yeast. Cell Cycle. 12:3228–3236. doi:10.4161/cc.26462.

Young, P.G., and P.A. Fantes. 1987. Schizosaccharomyces pombe mutants affected in their division response to starvation. J Cell Sci. 88 (Pt 3):295–304.

Zapata, J., N.E. Dephoure, T. MacDonough, Y. Yu, E.J. Parnell, M. Mooring, S.P. Gygi, D.J. Stillman, and D.R. Kellogg. 2014. PP2ARts1 is a master regulator of pathways that control cell size. J. Cell Biol. 204:359–376. doi:10.1083/jcb.201309119.

